# Algae Facade Technology in Improving Air Quality and Building Health: Analysis of Microclimate and Growth of *Aspergillus niger*

**DOI:** 10.1101/2023.02.01.526567

**Authors:** Muhammad Haikal Algifari, Ridzik Malky Daniel, Nofdianto

**Affiliations:** SMA Sukma Bangsa Lhokseumawe, Research Center for Limnology, National Research and Innovation Agency (BRIN); Research Center for Limnology, National Research and Innovation Agency (BRIN)

**Keywords:** Facade, *Chlorella pyrenoidosa*, *Aspergillus niger*, microclimate, photobioreactor

## Abstract

This research was conducted in response to concerns about room comfort and the Sick Building Syndrome (SBS) phenomenon. The focus of this study was to examine the effect of photobioreactor (PBR) based algae facade on the room microclimate and growth of *Aspergillus niger*. A glass simulation chamber resembling the letter “U” with a size of 40 cm long, 40 cm wide, and 40 cm high has been used for the experiment. Twenty samples of *Aspergillus niger* were tested for 11 days in four simulation chambers and were given different treatments, two rooms were installed with PBR and the other two rooms were not installed PBR (control). The room with PBR had a relatively lower temperature and humidity in the peak phase with a difference of 4.3°C and 5.5% RH compared to the control room. In addition, three out of ten molds in the PBR chamber had a growth rate of about 0.94 cm^2^/day and the spore production of 1.9×10^4^ CFU/ml. Meanwhile, seven out of ten molds in the control room had a growth rate of 1.4 cm^2^/day and an average spore production of 5.4×10^4^ CFU/ml. These results indicate that the algae facade is able to improve the microclimate of the room and suppress the growth rate of *Aspergillus niger*. mold in buildings.

## Introduction

In an effort to overcome the negative effects of unsustainable development practices and reduce their impact on an increasingly fragile natural environment, exploring biological systems and processes is gaining increasing attention. The integration of building facade microalgae is one of the inspirations in answering the above problems. Algae photobioreactor (PBR) is a bioreactor that utilizes sunlight as an energy source in photosynthesis to produce biomass. Building an algal symbiosis provides many benefits such as absorption of CO_2_ gas in the air, producing O_2_ which can improve air quality, and producing biomass (biofuel) for building energy [1], [2].

Apart from ensuring the energy issue, the algal facade also has a sun-shading effect and can effectively limit the heat in the building, so it will improve the quality of the indoor microclimate [1], [3]. This effect occurs because microalgae cells absorb the abiotic components of the sun for photosynthesis [4]–[6]. Another study controlling the temperature that is too high between outdoor and indoor on devices installed with algae PBR was presented by [7]. Meanwhile, [8] tested the symbiosis of algae and buildings to control room temperature in summer and winter.

On the other hand, microclimate problems frequently interfere with the comfort and health of the occupants as well as lead to the Sick Building Syndrome (SBS) phenomenon. SBS phenomenon commonly caused by the poor quality of temperature, humidity, and by the activity of spores, mycotoxins, and volatile compounds from toxic microorganisms such as fungus, bacteria, and other pathogenic organisms [9], [10]. However, many studies discussed that the microorganism, especially mold occurrence, is related to room microclimate [9], [11], [12].

One of the fungi commonly found in buildings is *Aspergillus sp*. [13]. This fungus highly not recommended to be indoors because it produces mycotoxins in the form of Aflatoxin B1, Fumonism, Gliotoxin, and Ochratoxin A [10], [14]–[17]. These mycotoxins carried on the spores of fungi which fly even under normal air ventilation. These mycotoxins could cause respiratory symptoms, allergies, and asthma if inhaled [15], [18]. In addition, this species of fungus also releases harmful Microbial Volatile Compounds (MVOC) such as alcohol, terpenes, etc. [10].

This study aims to measure the potential of algae facades in overcoming building health problems, especially the microclimate and growth of *Aspergillus niger* which often appears in buildings [16]. The results of this study are expected to contribute to stakeholders in solving energy and building health problems.

## Material and Methods

**Figure 1.**
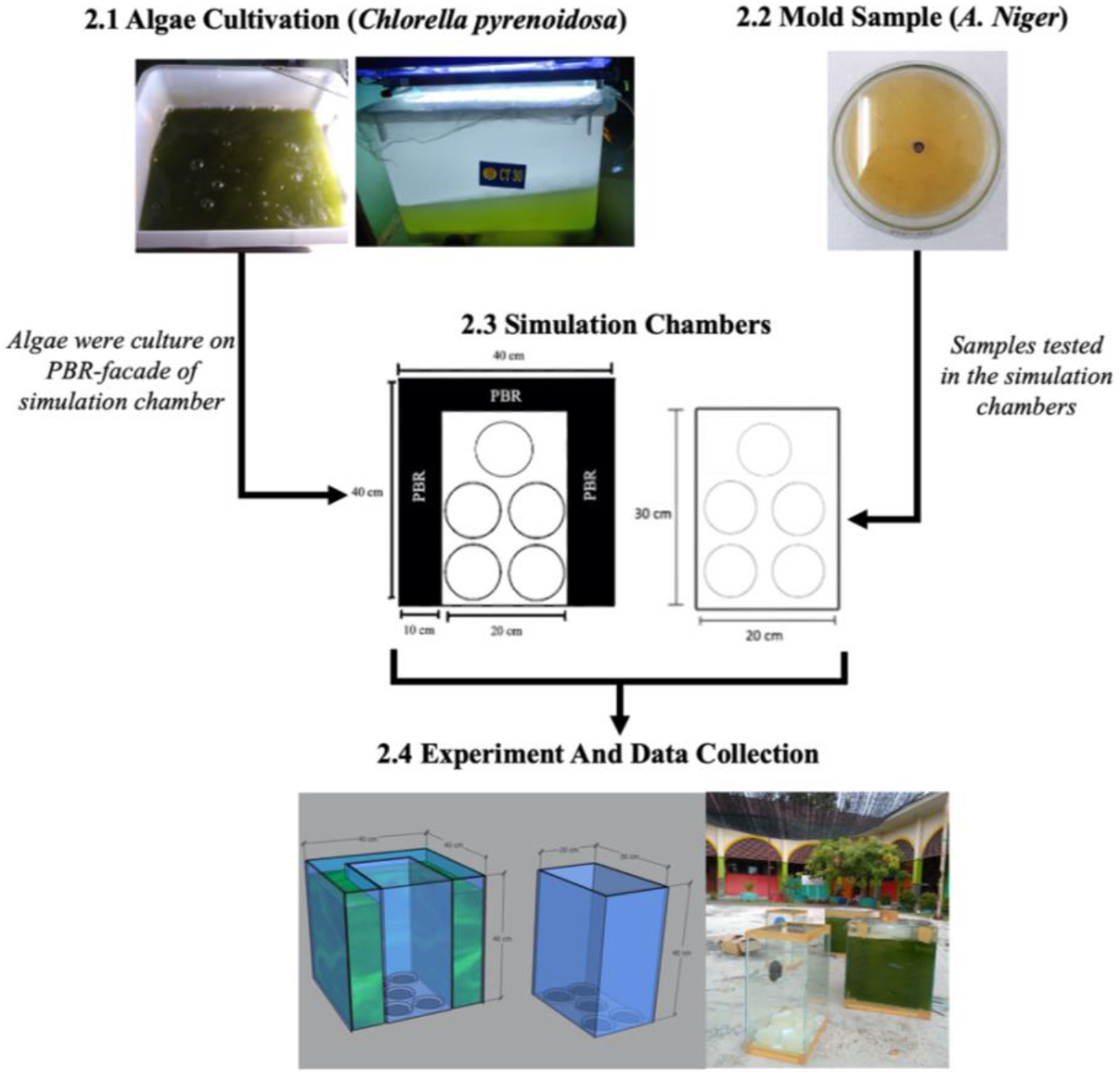
Flow of experimental procedure. The Algae *Chlorella pyrenoidosa* was cultivated inside PBR. Then *Aspergillus niger* was isolated to 20 Petri dish containing 4.8% MEA and tested under chamber’s microclimate. During the experiment, two simulation chambers were used [chambers using PBR as façade and without PBR (control)]. Then, the chambers were tested on real climate for 11 days and data collected by following: algae cell density, chambers and outdoor microclimate (temperature, humidity and light intensity), and mold growth area per day.

### 2.1 Algae Cultivation

Microalgae *Chlorella sp.* were commonly used in algae facade technology due to its durability, fast growth and high production of oxygen as well as absorbing carbon dioxide [19]–[22]. In this research, a *Chlorella pyrenoidosa* were used because its optimal growth were close to the real climate. [23] shows the optimal growth for *Chlorella pyrenoidosa* are 30°C which consider by the high production of biomass and lipid.

In the experiment, the seed of *Chlorella pyrenoidosa* (1 L) was purchased from BettaFish City, Bogor. Then, the seed was cultured (for seven days) by adding 5 L of distilled water and 500 mL fertilizers for nutrition. Fertilizer was made by diluting 30 g of bran, 10 mL of molasses, 18 mL of Effective Microorganism 1201179 (Persada, Jakarta), 4.30 g of urea 2801 (Pupuk Indonesia, Jakarta), and 30 g of NPK 2803 (Saprotan Utama, Semarang). Lastly, artificial light were added using Philips TL LED 16 W and aquarium air pump AA-350 AMARA were used to avoid cell deposition [24].

### 2.2 Mold Samples

To test the mold growth under the environment of simulation chambers, a total of 20 *A. niger* 6127 (FNCC, Yogyakarta) samples were prepared. The original culture of *A. niger* (approx. 80 cm^2^) was taken using a cork borer and placed into 20 new media on a petri dish. The petri dish contains 10 mL of Malt Extract Agar 1.05398.0500 (MEA) (Merck, Darmstadt). Agar plate was made by diluting 48 g of MEA into 1000 mL distilled water (4.8%). Lastly, the petri dish containing culture was sealed and readily tested on the chambers.

### 2.3 Simulation Chambers

Simulation of the algae facade building in the form of the letter “U” made of glass with a thickness of 5 mm with a total size of 40 cm long, 20 cm wide, and 40 cm high (Fig. 2). The “U” letter column with a volume of about 30 liters is a PBR facility for the algae *Chlorella pyrenoidosa* equipped with an AA-1880 aerator as a stirrer, and the space in the middle of about 0.024 cubic meters is a test site for microclimate and growth of *Aspergillus niger* mold. Meanwhile, the room without algae PBR installation was used as a control.

**Figure 2.**
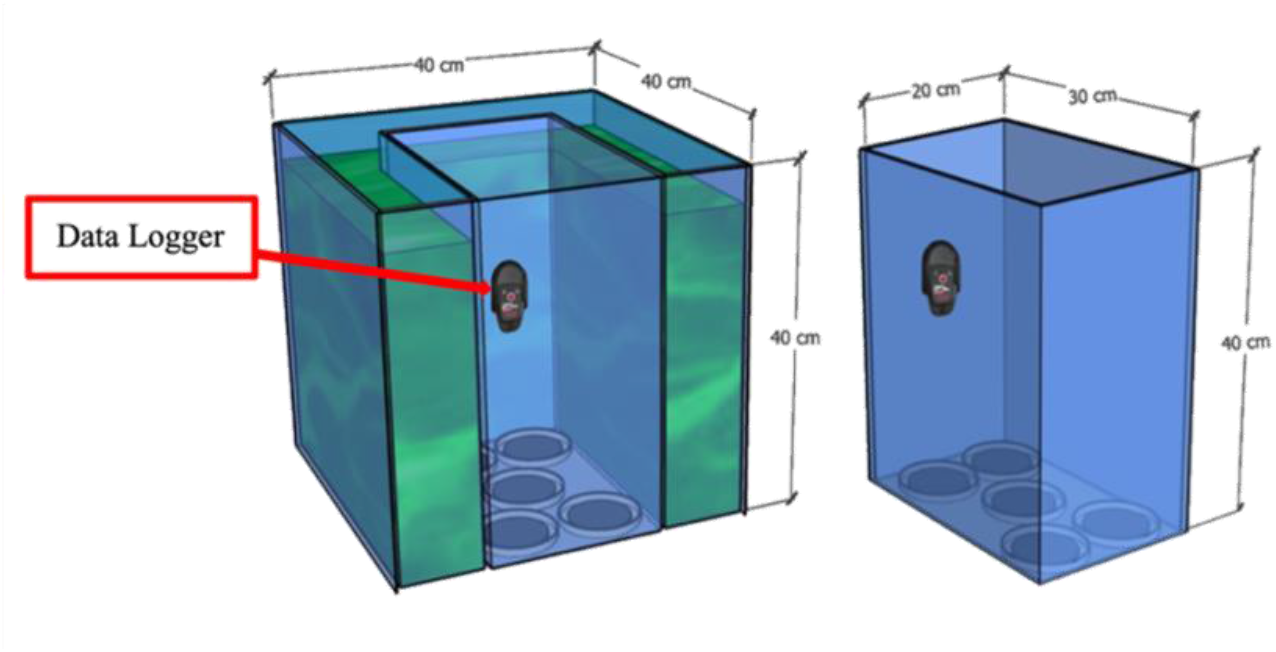
Experiment Devices: PBR device and control device

The simulation was carried out in an open space around the laboratory which was quite far from the resident’s settlements. A total of five petri dishes of *A. niger* mold were placed in the test room and in the control room, respectively. To avoid extreme weather, the test site is protected by 6×4 m of agricultural paranet.

### 2.4 Experiment and Data Collection

The experiment was carried by testing two types of simulation chambers in the climate of Sukma Bangsa High School, Aceh, Indonesia, for 11 days (started from October 10th – 20th 2020). In this experiment, the PBR-facade installation were the independent variables that affect the chamber’s microclimate (dependent variables). However, in the case of mold growth, the chamber microclimate could be an independent variables for the mold growth [14], [25]. Moreover, the control variables such as outdoor climate, the volume of agar, petri dish, and other materials are set as the same in this experiment.

The experimental data collected from experiment were chambers microclimate, algae cell density and mold growth area. For chambers microclimate, the USB-Data Logger BTH-81 placed inside the test chamber to record the temperature and humidity (Fig. 2). Data Logger was set to record the temperature and humidity every 3 minute during 11 days of experiment. Moreover, to measure the light that penetrated through the PBR, light intensity in the chambers was measured manually by Digital Lux Meter.

For algae measurements, cell density (cell/mL) was determined using a hemocytometer (Improved Neubauer). Approx. of 10 mL algae sample from each PBR will dilute with distilled water by dilution of 10^-3^ until 10^-4^ then the cell will be measured following by:

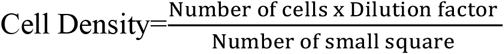

Lastly, for the mold growth area, a picture of 20 mold samples was taken one by one to calculate the growth area per day (specific growth rate). Image was calculated using an ImageJ blended with Java 1.8.0_172 software due to the high accuracy of measurement and less error. The mold picture of each day was imported to ImageJ software and the diameter of the petri dish was used to calibrate the scale (pixel to centimeters). Then, the outer of the petri dish was cut for clear measurement. The color threshold tools were used to distinguish the images based on the color difference. After the calculated sample was selected and marked by the yellow line, the “measure” button was clicked for the measurement result (Fig. 3).

**Figure 3.**
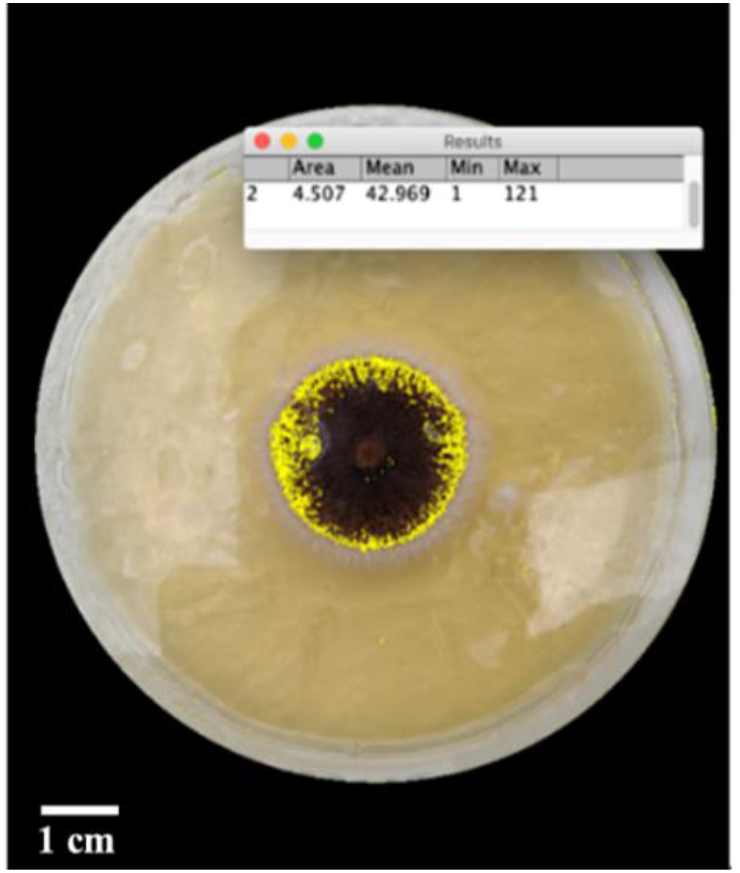
Measurement of growth area (cm^2^) using ImageJ Software.

### 2.5 Statistical analysis

To identify the differences between each dataset and treatment, Mann-Whitney U test and independent T test were applied in this experiment [28]. For the microclimate (temperature and humidity), the recorded data were not normally distributed, so a non-parametric test (Mann-Whitney U) was used. Meanwhile, for the growth of mold statistical testing using Parametric Test, independent T-test.

## Results And Discussion

### 3.1 Temperature

In Figure 4, the daily temperature during 11 days of the experiment was showed (Figure 4a). The temperature in the PBR chamber had a higher value compared to the control chamber at the night, especially when the sunlight begins to set (4.00 p.m. – 7.00 a.m.) and had a lower value at 10.00 a.m. – 4.00 p.m. or when the sunlight gradually increase. This result is alike [7] that mentioned the algae-PBR could decrease the indoor’s temperature when the outside temperature was lower. He also indicated algae-PBR could increase the temperature when the outside was low by releasing heat from algae-PBR that was exposed by solar radiation during the daytime [7].

**Figure 4.**
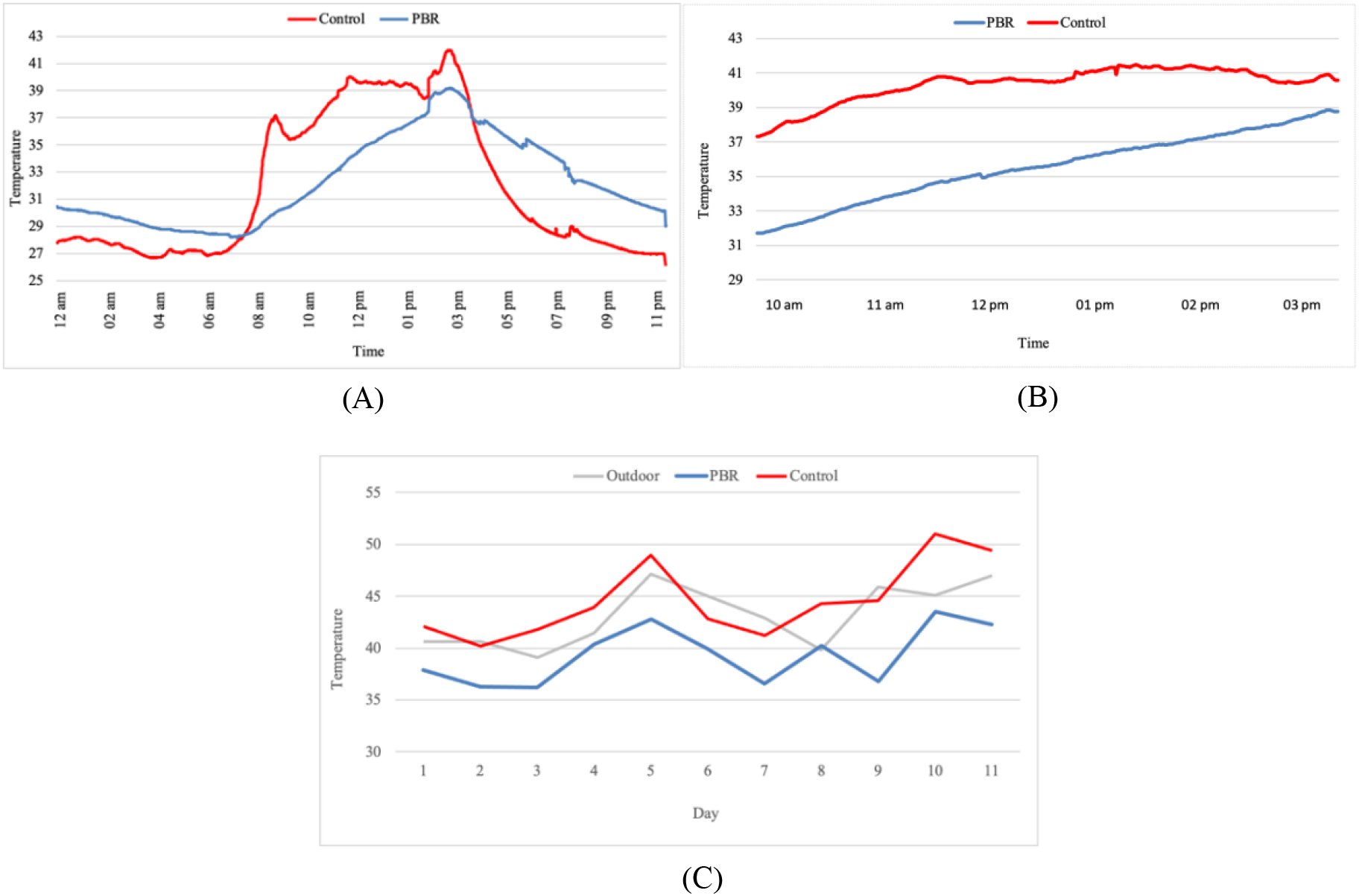
The temperature of the chambers (°C): (A) Daily Temperature on average, (B) Temperature during peak phase, (C) The Maximum temperature each chamber

The result also indicated that PBR installation could stabilize the chamber at the peak of outside temperature by a difference of 4.3°C (Figure 4b). This difference is highly related to the ability of PBR in absorbing high solar radiation. [3] Ismail and Al Obaidi (2019) mentioned that PBR panel could prevent up to 44.9% of solar radiation and 25% of solar transmission that possibly reduce heat gain of the building. Figure 4c also shows the maximum value recorded on PBR chambers was lower than the control chamber by difference of 5.3°C on average (Figure 4c).

### 3.2 Humidity

Fig. 5a shows that from 8.00 a.m. to 4.00 p.m. the humidity in the control chamber was lower than PBR chamber by a difference of 16.9% RH. While at 06:00 pm to 08:00 am, the humidity in the control chamber was higher than in the PBR chamber by a difference of 4.8%RH (Fig. 5a). This humidity value is strongly related to temperature [14]. According to the Figure 4a and 5a, the correlation between temperature and humidity are shown. When the temperature was higher (8.00 a.m. to 4.00 p.m.), the humidity was lower, and *vice versa* (Figure 4a and 5a). The humidity could decrease drastically due to the excessive heat that could reduce the water activity in the air by evaporating process.

**Figure 5.**
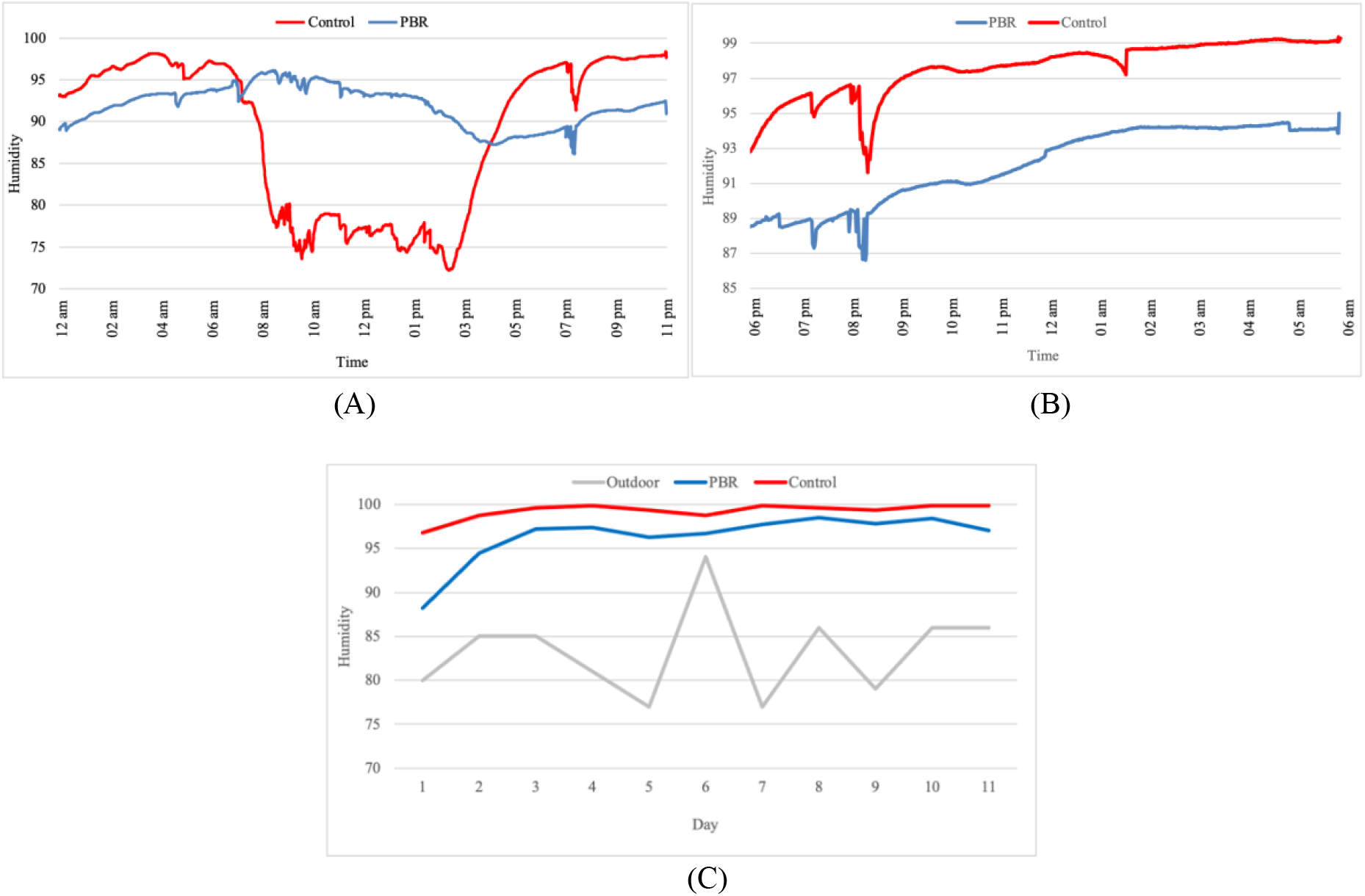
The humidity of the chambers (%RH): (A) Daily humidity on average, (B) Humidity during peak phase, (C) The Maximum humidity of each chamber

Furthermore, when the outside humidity was higher due to the absence of heat from the sun or called peak phase (6.00 p.m. to 8.00 a.m.), the PBR chambers had a lower humidity value than the control chamber (Figure 5b). This finding is related to the heat that is released by PBR during the low temperature of the outside [7]. Furthermore, the heat exposed by algae to the chamber will determine the humidity value since the correlation between temperature-humidity is already stated [14].

### 3.3 Light Intensity

In the PBR chamber, the light acceptance rate is 2.295 Lux on average or 11.2% while in the control chamber is 11.044 Lux or 51.8% (Fig 4d). The low penetration of light in the PBR chamber was caused by microalgae cell density that absorbs heat and radiation from the sun [3]. Also, PBR could prevent the sun from direct exposure or is called the *sun-shading* effect [1]. These shading and prevention systems could be beneficial for the heat gain of buildings, including provide a natural cooling system and improving thermal comfort [2], [7].

**Figure 6.**
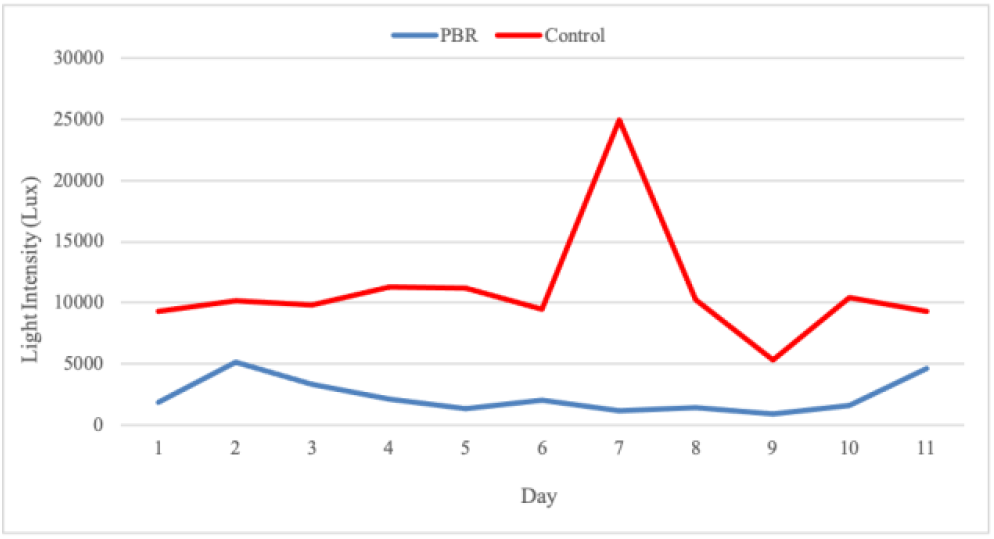
The Light acceptance of the chambers (Lux)

### 3.4 Algae Cell Density

Algae cell growth during the experiment ranged from 50×10^4^ to 140×10^4^ cells/ml (Fig. 7), and algae still could grow very well signed by the changes of the algae color to the deep dark green [1]. Cell density certainly affects the quality of the shade and the surrounding temperature. However, the installation of algae facades in the sun for a long time can cause growth inhibition effects and cell damage [26]. According to [27], north or south orientation is better at latitudes below 35°C to prevent direct sun exposure.

**Figure 7.**
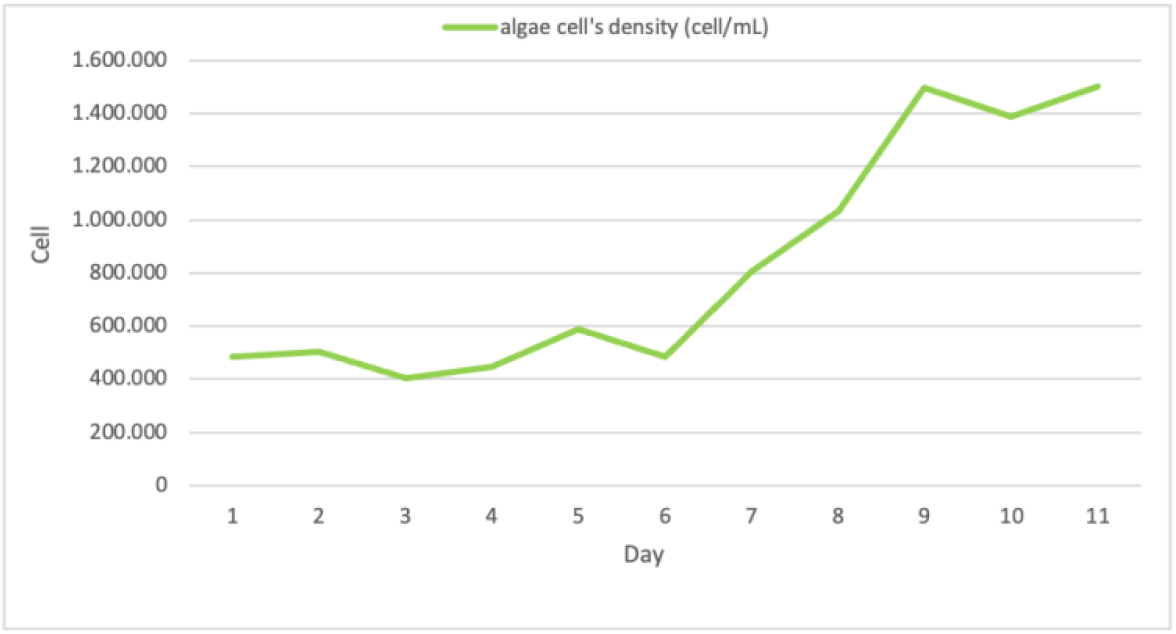
Algae cell density (cell/mL)

### 3.5 Mold Growth

#### (a) Fungal occurrence and mold features

A total of 20 samples were tested, only ten samples were observed for growth and the rest indicated contaminants. The total growth of *A. niger* fungus on the first day was higher in the control room than in the PBR room. These results indicate that there are differences in environmental conditions and microclimate between the two test rooms. Some literatures say that the growth and germination of fungi is not only influenced by the substrate, but also by temperature and humidity [25]. In this case, the PBR and control test chambers gave an average humidity of 87.8% and 96.1% RH, temperatures of 30.6°C and 27.1°C, respectively. The PBR room showed a negative effect on the growth of *A. niger*, and the control room not only supported the growth of *A. niger* species with humidity close to 98%RH [14], also for the other three species that germinated on 5^th^ and 8^th^ day.

The morphological characteristics of fungi with PBR and control treatments (Table 1) also showed differences. In the PBR room, there were few black spores, and white mycelium. Meanwhile, in the control room, there were more black spores, the mycelium was very thin and almost invisible because it was covered by spores. This difference indicates that temperature and humidity are essential factors affecting the production phase of fungal spores and conidia [14], [25]. In its life cycle, the spores will germinate, if the substrate has provided sufficient moisture and nutrients. After the spores germinate, hyphae (fungal filaments) are formed, and continue to grow to form lateral branches. With sufficient humidity conditions, the hyphae will thicken and form a mycelium. At this stage, the fungus metabolizes and retains sufficient moisture to sustain growth [25].

**Table 1.**
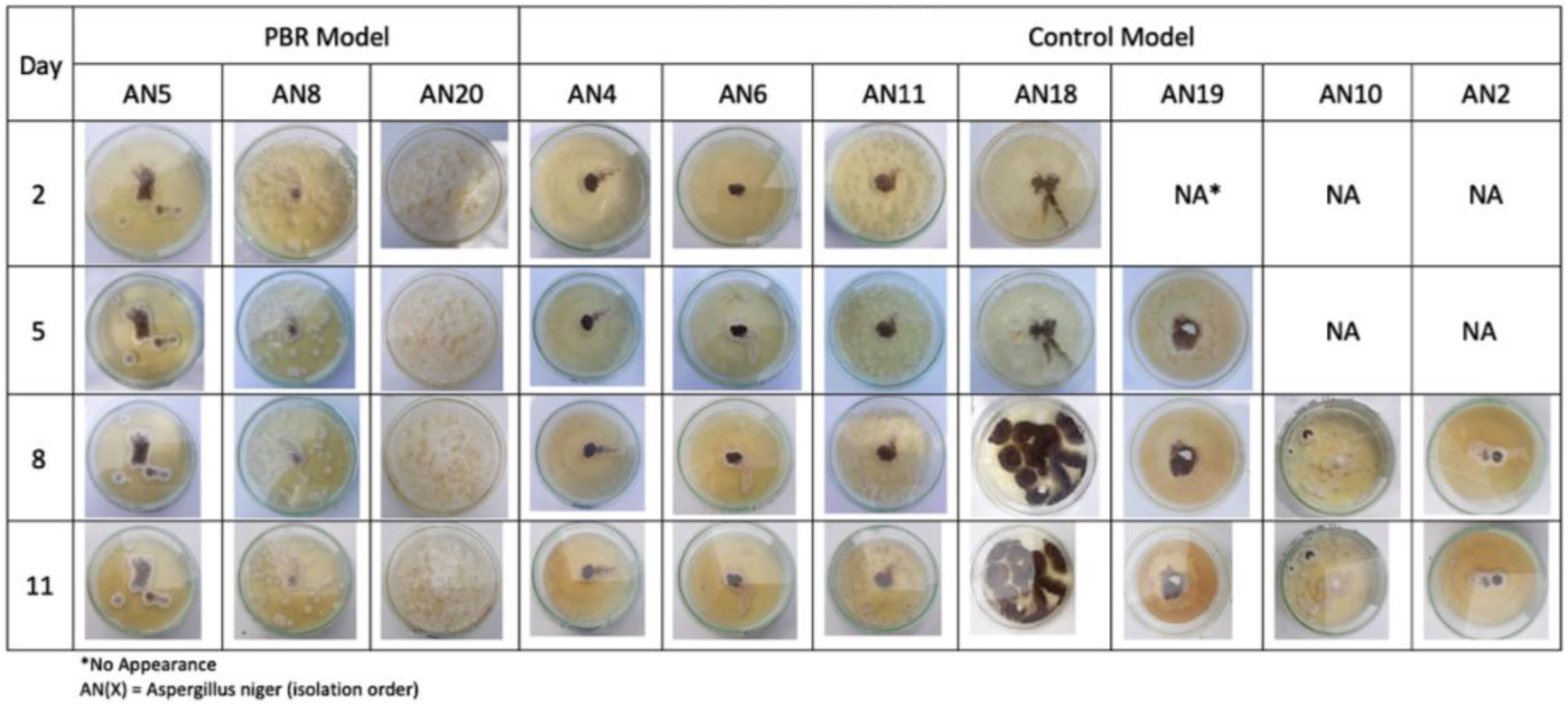
Mold (*Aspergillus niger*) appearance.

#### (b) Total Area and Conidia

Furthermore, the difference in growing area (cm^2^/day) in the PBR room was lower than in the control room (Fig. 5). In the PBR sample, the mold grew on average 0.94 cm^2^/day, whereas in the control room it grew 1.41 cm^2^/day. In the conidia production (cfu/mL), the PBR room had an average conidia of 1.9×10^4^ cfu/mL, while the sample in the control room was 5.4×10^4^ cfu/mL. This parameter also proves that the mold samples in the PBR room experienced obstacles in the process of reproduction and growth compared to the control room.

**Figure 5.**
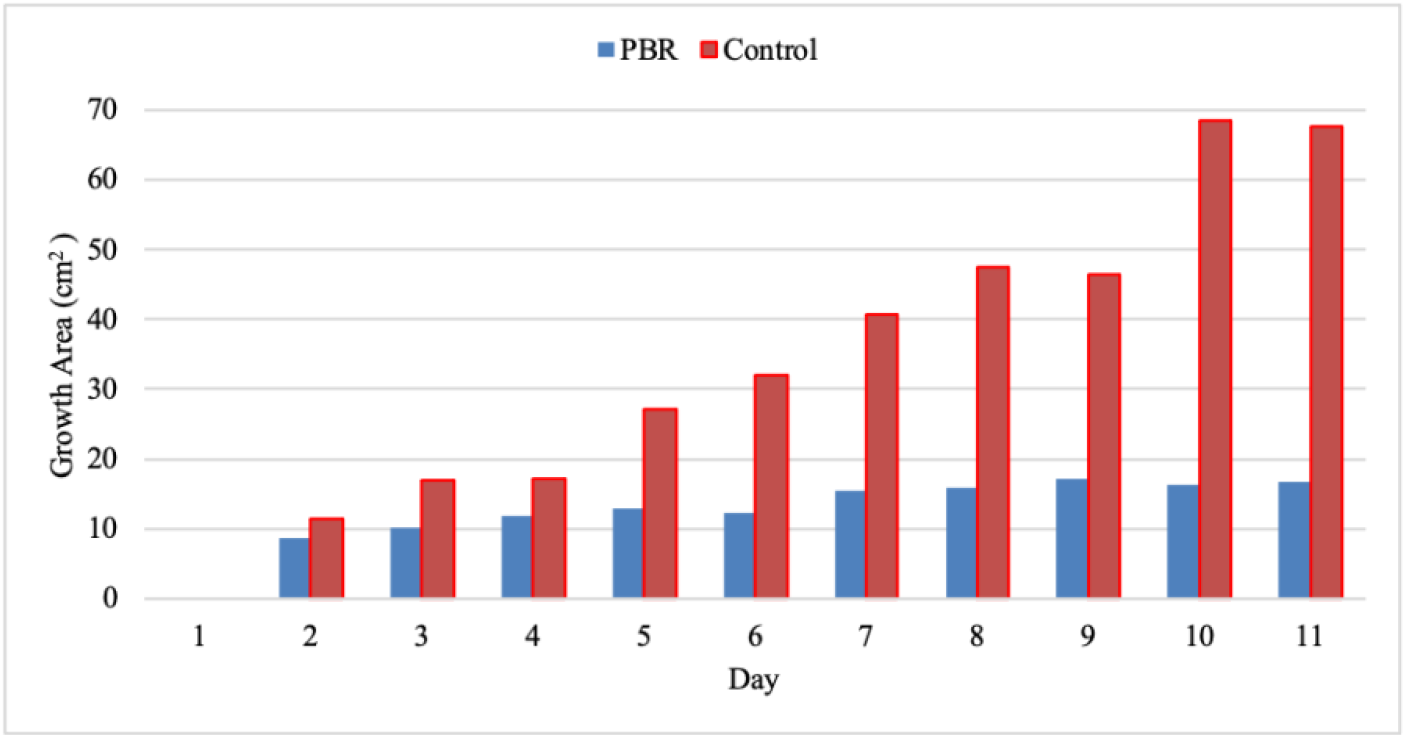
Total mold area in PBR and control samples (cm^2^)

### 3.6 Statistical Test

The results of statistical tests showed that the microclimate (temperature and humidity) between the algae facade and the control room was significantly different < 0.005 (Fig. 6a, b). Similar results also occurred in the area of fungal growth in both test chambers (Fig. 6c).

**Figure 6.**
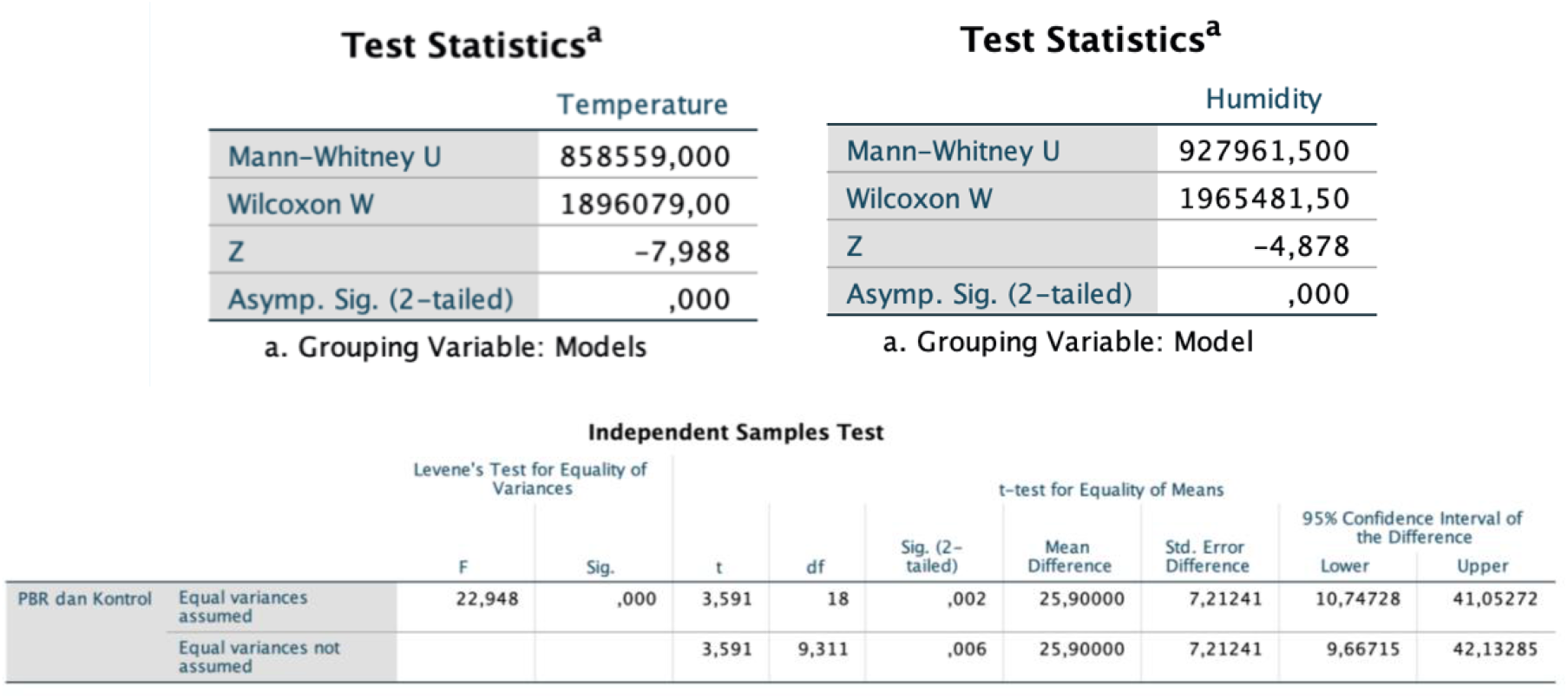
(a) Mann-Whitney test for temperature. (b) Mann-Whitney test for humidity. (c) Independent T test for mold growth area.

## Conclusion

Several points that can be underlined from the results of this study are: 1. Installation of algal facades has the potential to improve the microclimate and control the growth rate of *Aspergillus niger* mold in a building; 2. The algal facade provides coolness by lowering the temperature during the day and humidity at night; 3. Weather factors, especially in the tropics, may affect the results of this study, especially the duration of the rainy and hot seasons.

## Perspective

Due to the previous research and the findings in this study suggest a close relationship for environmental microclimate related mold growth, the installation of this algae facade has the potential to impact other microorganisms in the building too (such as pests, insects, and other pathogenic organisms such as viruses or bacteria). However, the improvements in methodology and variable analysis should be considered in future studies (e.g., MVOC or mycotoxin production, organism/pest behavior, etc.). In addition, real climate testing in one year is required, as well as a better simulation or computer simulation tool (ESPr, EnergyPlus).

## Acknowledgements

This research is the winner of the Youth Scientific Paper Competition (LKIR) held by the Indonesian Institute of Sciences and participated in the scientific competition at the International Science and Engineering Fair (ISEF) 2021. Thanks to teacher Fauza Azima, mentor, and the Food and Microbiology Laboratory, Chemical Engineering, Lhokseumawe State Polytechnic which has facilitated this activity.

